# Modulation of dysbiotic vaginal complications by cervical mucus revealed in linked human vagina and cervix chips

**DOI:** 10.1101/2023.11.22.568273

**Authors:** Ola Gutzeit, Aakanksha Gulati, Zohreh Izadifar, Anna Stejskalova, Hassan Rhbiny, Justin Cotton, Bogdan Budnik, Sanjid Shahriar, Girija Goyal, Abidemi Junaid, Donald E. Ingber

## Abstract

**Background:** The cervicovaginal mucus that coats the upper surface of the vaginal epithelium is thought to serve as a selective barrier that helps to clear pathogens, however, its role in modulating the physiology and pathophysiology of the human vagina is poorly understood. Bacterial vaginosis (BV), a common disease of the female reproductive tract that increases susceptibility to sexually transmitted infections, pelvic inflammatory disease, infertility, preterm birth, and both maternal and neonatal infections is characterized by the presence of a wide array of strict and facultative anaerobes, often including *Gardnerella vaginalis*.

**Objective:** To assess the role of cervical mucus in preventing dysbiosis-associated complications and preserving vaginal health.

**Study Design:** To better understand the role of cervicovaginal mucus in vaginal health, we used human organ-on-a-chip (Organ Chip) microfluidic culture technology to analyze the effects of cervical mucus produced in a human Cervix Chip when transferred to a human Vagina Chip BV model. Both chips are lined by primary human organ-specific (cervical or vaginal) epithelium interfaced with organ-specific stromal fibroblasts.

**Results:** Our data show that mucus-containing effluents from Cervix Chips protect Vagina Chips from inflammation and epithelial cell injury caused by co-culture with dysbiotic microbiome containing *G. vaginalis*. Proteomic analysis of proteins produced by the Vagina Chip following treatment with the Cervix Chip mucus also revealed a collection of differentially abundant proteins that may contribute to the vaginal response to dysbiotic microbiome, which could represent potential diagnostic biomarkers or therapeutic targets for management of BV.

**Conclusions:** This study highlights the importance of cervical mucus in control of human vaginal physiology and pathophysiology, and demonstrates the potential value of Organ Chip technology for studies focused on health and diseases of the female reproductive tract.

## INTRODUCTION

The cervicovaginal fluid that covers the surface of the vaginal epithelium is essential to women’s health and reproductive functions because it serves as a selective barrier that protects against environmental pathogens.^1, 2^ This fluid contains mucus that is mainly produced by the cervical epithelium along with vaginal secretions, and it contains a range of different cytokines, chemokines, immunoglobulins, and other immune mediators that help to prevent pathogens from crossing this critical mucosal interface. However, the role of the cervical mucus in regulating vaginal microbiome composition and its effects on health outcomes has largely remained unexplored.^3^ This is important because dysbiotic changes in the composition of the female genital tract microbiome, as observed for example in patients with bacterial vaginosis (BV) whose microbiome often contains strict and facultative anaerobes including *Garderenella vaginalis*^4, 5^ have been linked to increased susceptibility to sexually transmitted infections, pelvic inflammatory disease, infertility, preterm birth, and maternal and neonatal infections.^6, 7^ BV is also the most prevalent cause of vaginal symptoms in women with a more than a 50% recurrence rate, yet the underlying factors contributing to these conditions remain elusive.^8^

In this study, we leveraged human organ-on-a-chip (Organ Chip) microfluidic culture technology to explore directly how cervical mucus secretions influence human vaginal epithelium under both healthy and dysbiotic conditions. To do this, we connected two recently described human Organ Chip models of female reproductive organs. The first is a human vagina-on-a-chip (Vagina Chip) that is lined by primary, hormone-sensitive, vaginal epithelium interfaced with underlying stromal fibroblasts that we have shown recapitulates the pathophysiology of a dysbiotic vaginal epithelium when co-cultured with a *G. vaginalis* containing microbiome and that enables analysis of human host-microbiome interactions in vitro.^9^ The second is a human Cervix Chip lined by primary cervical epithelium interfaced with cervical fibroblasts^10^ that produces abundant cervical mucus with compositional, biophysical, and hormone-responsive properties similar to those observed *in vivo*. To model and study the effect of cervical mucus on vaginal responses in vitro, we co-cultured dysbiotic microbiome in the human Vagina Chip in the presence or absence of mucus-containing effluents that were transferred from the epithelial channel of the human Cervix Chips. These studies revealed that human cervical epithelial secretions exert immunomodulatory effects and protect the vaginal epithelium against a dysbiotic microbiome by reducing innate inflammatory responses and inhibiting growth of *G. vaginalis* bacteria, thereby reducing vaginal cell injury.

## MATERIALS AND METHODS

### Human Vagina Chip Culture

The Human Vagina Chip was cultured as previously described (36434666). In summary, microfluidic two-channel co-culture Organ Chip devices (CHIP-S1TM) were obtained from Emulate Inc. (Boston, MA). The Polydimethylsiloxane (PDMS) membrane was coated with collagen IV (30 μg/mL) (Sigma, cat. no. C7521) and collagen I (200 μg/mL) (Corning, cat. no. 354236) in Dulbecco’s Modified Eagle Medium (DMEM, ThermoFisher, cat. no. 12320-032) in the apical channel. The basal channel was coated with collagen I (200 μg/mL) (Corning, USA) and poly-L-lysine (15 μg/mL) (ScienCell, Cat# 0403).

Primary human uterine fibroblasts (ScienCell Research Laboratories, cat. no. 7040) were then seeded at a density of 1 × 10^6^ cells/mL in the basal channel and the human vaginal epithelial cells (Lifeline Cell Technology, cat. no. FC-0083; donors 05328) were seeded at a density of 3 × 106 cells/mL in the apical channel. The chips were incubated at 37°C with 5% CO2 under static conditions till the cells formed a uniform monolayer. The chips were then connected to the culture module instrument (ZOË™ CULTURE MODULE, Emulate Inc., USA) and put under flow conditions. The Vagina Chips were cultured using a periodic flow regimen in which vaginal epithelium growth medium (Lifeline, Cat# LL-0068) was flowed through the apical channel for four hours per day at 15 μl/hr. The basal channel was flowed continuously with fibroblast growth medium (ScienCell, Cat# 2301) at 30 μl/hr. After 5-6 days, the basal medium was replaced with an in-house differentiation medium^9^ for eighth days following the same intermittent and continuous perfusion regime in the apical and the basal channels respectively. The apical medium was replaced with customized HBSS Low Buffer/+Glucose (HBSS (LB/+G)) and the basal medium was replaced with antibiotic free differentiation medium for one day followed by three days with microbial co-culture as described below.

### Human Cervix Chip Culture

Human Cervix Chip Culture was cultured as previously described (Izadifar et al., 2023, BioRxiv). In summary, microfluidic two-channel co-culture Organ Chip devices (CHIP-S1^TM^) were obtained from Emulate Inc. (Boston, MA). The PDMS membrane was coated with 500 μg/ml collagen IV (Sigma-Aldrich, Cat. no. C7521) in the apical channel and with 200 μg/ml Collagen I (Advanced BioMatrix, Cat. no. 5005) and 30 μg/ml fibronectin (Corning, Cat. no. 356008) in the basal channel. Primary cervical fibroblasts (0.65 x10^6^ cells/ml, P5)(isolated from hysterectomy cervical tissues) were seeded on the basal side followed by seeding the primary cervical epithelial cells (1.5 x 10^6^ cells/ml, P5) (LifeLine Cell Technology Cat# FC-0080) on the apical side. The respective media of the chips were refreshed after the seeding process for each channel and the chips were incubated at 37°C, 5% CO2 under static conditions overnight. The chips were then connected to the culture module instrument (ZOË™ CULTURE MODULE, Emulate Inc.,USA). The Cervix Chips were cultured using a periodic flow regimen in which cervical growth medium was flowed through the apical channel for four hours per day at 30 μl/hr while fibroblast growth medium was continuously perfused basally at 40 μl/hr. After five days the apical medium was replaced by Hank’s Buffer Saline Solution (HBSS) (Thermo Fisher, 14025076) while being fed through the basal channel by differentiation medium constitue of cervical epithelial medium (LifeLine Cell Technology, Cat. no. LL-0072) supplemented with 5 nM estradiol-17β (E2) (Sigma, Cat. no. E2257) and 50 µg/mL Ascorbic acid (ATCC, Cat. no. PCS-201-040) at day 2 of differentiation the apical medium was replaced by a customized HBSS with low buffering salts and no glucose (HBSS (LB/-G) with pH ∼5.4 and cultured for 5 additional days.

### Mucus collection from the cervix chip

Cervix Chip mucus was collected every day starting day 4 of differentiation for 10 days. During the collection period, the basal channel was continuously perfused with antibiotics free differentiation medium at a volumetric flow rate of 40 μL/h. The apical channel was perfused with HBSS (LB/-G) for 4 hours per day at 40 μL/h flow rate and the chip effluetns were collected and stored at -80°C till the time of the experiment. Before adding the mucus to the Vagina Chip 5.56 mM D-glucose (Sigma, cat. no. G7021) was added to the chip mucus.

### Culture of a non-optimal *Gardnerella vaginalis* containing Consortium in the Vagina Chip

In non-optimal vaginal microbiota, *Gardnerella* species are typically found as dominant bacteria^7^ accompanied by other frequent taxa such as *Prevotella* species and *Atopobium* species.^11^ To mimic the ecology of non-optimal vaginal microbiota, we used a synthetic dysbiotic consortia (BVC1: *Gardnerella vaginalis E2, Gardnerella vaginalis E4, Prevotella bivia BHK8, and Atopobium vaginae*). The *Gardnerella* isolates used in this study were selected because they represent distinct genomic groups, exhibit phenotypic diversity in vitro, and were co-resident, meaning that they were co-isolated from a single participant in the UMB-HMP study.^12^ *P. bivia* and *A. vaginae* are prevalent species in *Lactobacillus*-deficient vaginal microbiota. The two strains used in this study were co-resident, isolated from a single participant in the Females Rising Through Education Support and Health study.^13^ The apical chip channel was inoculated with ∼10^5^ CFU of prepared BVC1 consortia and then chips were incubated statically at 37°C and 5% CO2 for 20 hours before starting the flow using the Zoe culture module. The basal channel was continuously perfused with in-house antibiotic free differentiation medium and apical channel was perfused for 4 hours per day with customized HBSS (LB/+G) medium at a volumetric flow rate of 40 μL/hr.

### Study design

The study was carried out using five conditions: control, BVC1, pre-treatment, post-treatment and pre+post treatment. Vaginal epithelium cultured on-chip for 72 hours in the absence (Control) or presence of BVC1 consortium either without or with mucus. Mucuse pre-treatment group, where cervical mucus was used as the apical medium for 24 hours before BVC1 infection followed by the use of customized HBSS Low Buffer/+Glucose (HBSS (LB/+G)) as the apical medium. Mucus post-treatment group, where cervical mucus was used as the apical medium for the duration of the experiment, beginning 24 hours after BVC1 infection. Mucus pre+post treatment group, where cervical mucus was used as the apical medium for 24 hours prior to BVC1 infection and for 72 hours during BVC1 co-culture.

### Bacterial Enumeration from Vagina Chip Co-Culture

To enumerate all cultivable bacteria in the effluents, effluent samples (50 μL) were collected at 24, 48, and 72 hours. Effluent samples from the Vagina Chips containing BVC1 consortia were plated on Brucella blood agar (with hemin and vitamin K1) (Hardy, cat. no. A30) at 37°C under completely anaerobic conditions. After 48 hours of incubation, colonies were counted, and CFU/chip was calculated for each sample. To enumerate all cultivable bacteria engrafted in the Vagina Chip, the whole cell layer was digested for 1 hour with a digestion solution containing 1 mg/mL of collagenase IV (Gibco, cat. no. 17104019) in TrypLE (ThermoFisher, cat. no. 12605010). Cell layer digest was then diluted and processed in the same way as effluent samples and CFU/chip was calculated for each chip digest.

### Analysis of Cytokines and Chemokines

Samples (100 μL) of the apical effluents from Vagina Chips were collected and analyzed for a panel of cytokines and chemokines, including TNF-α, IFN-y, IL-1α, IL-1β, IL-10, IL-8, IL-6, MIP-1α, MIP-1β, IP-10, and RANTES using custom ProcartaPlex assay kits (ThermoFisher Scientific). The analyte concentrations were determined using a Luminex 100/200 Flexmap3D instrument coupled with the Luminex XPONENT software.

### Protein extraction and mass spectrometry

Digestion of the samples: Samples were run through 50 kDa filter (Amicron Ultracel, Merck Millipore, Ireland) and digested according to the manafacturer’s protocol for 1 hour at 50 °C by Trypsin Platinum (Promega, WV), digested material was dried in speedvac (Eppendorf, Germany). Mass spectrometry analysis: After digestion each sample was resolubilized in 10 ul of 0.1 % formic acid in water buffer A solution. Each sample submitted for single LC-MS/MS experiment that was performed on a 240 Exploris Orbitrap (ThermoScientific, Germany) equipped with NEO nano-HPLC pump (ThermoScientific, Germany). Peptides were separated onto a micropac 5 cm trapping column (Thermo, Belgium) followed by 50 cm micropac analytical column 50 cm of (ThermoScientific, Belgium). Separation was achieved through applying a gradient from 5–24% ACN in 0.1% formic acid over 90 min at 250 nl min−1. Electrospray ionization was enabled through applying a voltage of 1.8 kV using electrode junction (PepSep, Denmark) at the end of the microcapillary column and sprayed from stainless-steel 4 cm needle (ThermoScientific, Denmark). The Exploris Orbitrap was operated in data-dependent mode for the mass spectrometry methods. The mass spectrometry survey scan was performed in the Orbitrap in the range of 450 –1,200 m/z at a resolution of 1.2 × 105, followed by the selection of the ten most intense ions (TOP10) for HCD-MS2 fragmentation in the orbitrap. The fragment ion isolation width was set to 0.8 m/z, AGC was set to 50,000, the maximum ion time was 150 ms, normalized collision energy was set to 32V and an activation time of 1 ms for each HCD MS2 scan.

Data analysis: Raw data were submitted for analysis in Proteome Discoverer 3.0 (Thermo Scientific, CA) software. Assignment of MS/MS spectra was performed using the Sequest HT algorithm by searching the data against a protein sequence database including all entries from the Human Uniprot database (SwissProt, 2019) and full Uniprot bacteria database (SwissProt, 2022) as well as known contaminants such as human keratins and common lab contaminants. Sequest HT searches were performed using a 15 ppm precursor ion tolerance and requiring each peptides N-/C termini to adhere with Trypsin protease specificity, while allowing up to two missed cleavages. For searches methionine oxidation (+15.99492 Da) and asparagine and glutamine deamidations (+0.984016 Da) were set as variable modification as well as N-terminal acetylation of protein terminus. A MS2 spectra assignment false discovery rate (FDR) of 1% on protein level was achieved by applying the target-decoy database search. Filtering was performed using a Percolator (64bit version).^14^ For quantification analysis between samples label free quantitation mode using Minora detection features of Proteome Discoverer platform was used.

### Statistical analysis

All the results presented are from at least two independent experiments and all of the data points shown indicate the mean ± standard deviation (s.d.) from n > 3 Organ Chips unless otherwise mentioned. Tests for statistically significant differences between groups were performed using unpaired t-test, statistical analyses were performed using GraphPad Prism 9.0.2.

## RESULTS

### Modulation of innate immunity in Vagina Chips by mucus-containing effluents from Cervix Chips

We recently described a human Vagina Chip lined by primary human vaginal epithelium interfaced across an extracellular matrix (ECM)-coated porous membrane with underlying stromal fibroblasts cells that enables analysis of human host-microbiome interactions in the vaginal microenvironment,^9^ as well as a human Cervix Chip containing a primary cervical epithelium interfaced with stromal cervical fibroblasts that produces cervical mucus with physical and chemical properties similar to those observed *in vivo*.^10^ Here, we collected mucus-containing effluents from the outflow of the epithelial channel of the Cervix Chip (’cervical chip mucus’ containing 4.01 ± 3.04 mg/ml of mucus glycoproteins) for 7 days and then perfused it through the epithelial channel of a Vagina Chip to simulate the natural flow of mucus in the reproductive tract in vivo (**Fig. 1A**). The presence of this mucus in the Vagina Chip induced statistically significant decreases in secretion of multiple relevant proinflammatory cytokines, including interleukin-1α (IL-1α), IL-1β, and macrophage inflammatory protein-1β (MIP-1β), accompanied by a concomitant increase in anti-inflammatory IL-10 protein production after 24 hours of exposure compared to control chips without mucus (**Fig. 1B**). These results demonstrate that the mucus-containing fluids produced by human cervical epithelium in Cervix Chips in vitro can directly influence the vaginal epithelium and result in suppression of production of inflammatory cytokines, even in the absence of immune cells.

**Fig. 1.**
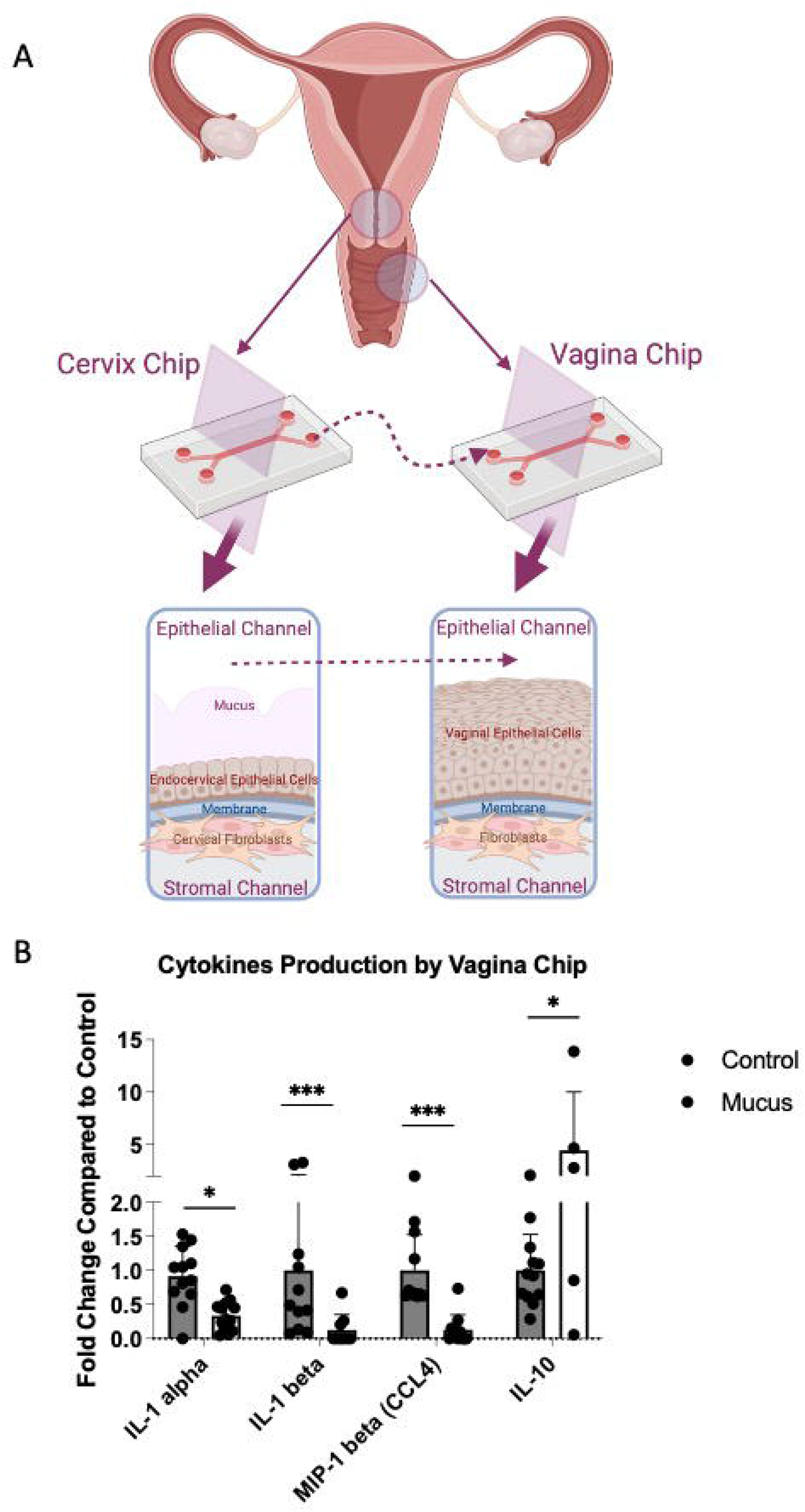
Effects of Cervical Mucus on Cytokine Production by the Vagina. A) Schematic diagrams of the Cervix and Vagina Chips and the transfer of cervical mucus between the chips. B) Cytokine protein levels for IL-1α, IL-1β, MIP-1β and IL-10 measured in effluents of Vagina Chips cultured with (light gray bars) without (dark gray bars) cervical mucus for 1 day. Each data point indicates one chip; data shown are from 3 different experiments and are presented as mean ± sd; significance was calculated by unpaired t-test; ***, P< 0.0001; **, P < 0.001.

### Modulation of the dysbiotic vaginal microbiome by introducing cervical mucus into the Vagina Chip

We next studied the effects of cervical mucus on a dysbiotic (non-optimal) vaginal microbiome in the Vagina Chip by inoculating the chip with a consortium containing G. *vaginalis E2* and *E4* combined with *P. bivia BHK8* and *A. vaginae* (BVC1; ∼10^5^ CFU/chip) on day 14 of culture in the presence or absence of mucus-containing effluents from the Cervix Chip. Interestingly, the presence of human cervical mucus inhibited the consortium’s ability to colonize the epithelium and thrive in the Vagina Chip. The total number of colony forming units (CFU) of live non-adherent bacteria collected in effluents from the epithelial channel during 72 hours of infection (**Fig. 2A**) as well as number of live adherent bacteria in tissue digests at the end of the 72 hour culture (**Fig. 2B**) were significantly reduced whether the Vagina Chips were pretreated with mucus effluents for 1 day before introduction of the microbiome, 1 day after BVC 1 addition, or continuously for the entire 3 day culture starting 1 day prior to addition of bacteria.

**Fig. 2.**
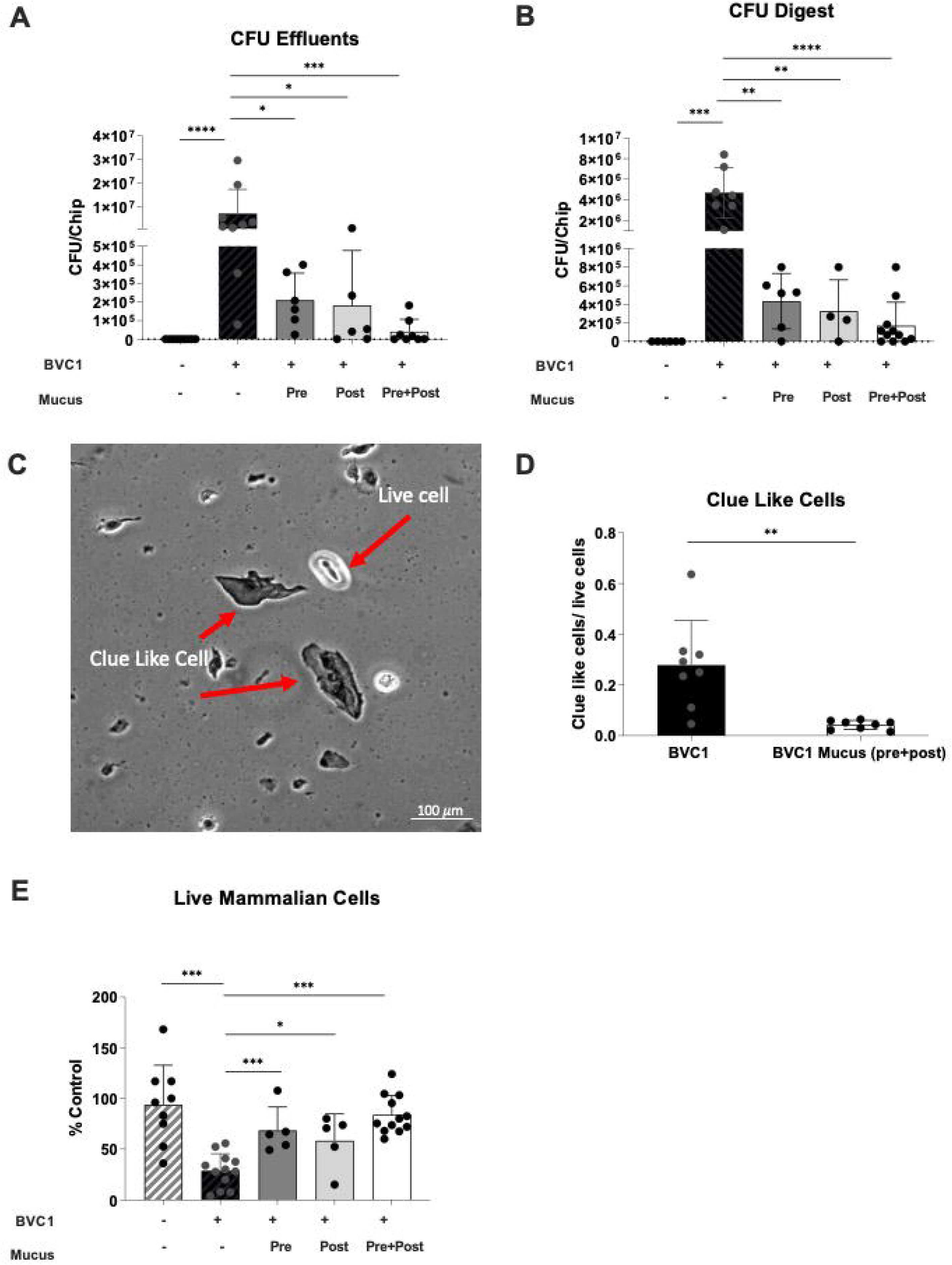
Modulation of dysbiotic vaginal microbiome by Cervical Mucus introduced into the Vagina Chip. Vaginal epithelium cultured on-chip for 72 hours in the absence (Control: grey with stripes bars) or presence of BVC1 consortium (black bars) perfused either without or with mucus-containing effluents from Cervix Chips that were added 1 day prior to addition of bacteria (Pre; dark grey bar), 1 day after BVC 1 addition (Post; light grey bar), or continuously for the entire 3 day culture starting 1 day prior to addition of bacteria (Pre+Post; white bar). **A**) Total non-adherent bacterial cell number (CFU) per chip determined by quantifying bacteria collected in effluents from the apical epithelial channel during 72 hours of co-culture of BVC1 in the Vagina Chip. **B**) Total adherent CFU per chip determined by quantifying of bacteria retained within epithelial tissue digests after 72 hours of culture. **C**) Quantification of vaginal epithelial cell injury (percent cell viability) assessed by calculating the number of live cells relative to control using Trypan blue exclusion assay. **D**) Bright field microsopic image showing Clue-like cells and live epithelial cells staining using Trypan blue. **E**) Ratio of Clue-like cells to live cells detected as described in **D**. In all graphs, results were obtained from at least 2 different experiments; each data point indicates one chip. Data are presented as mean ± sd; significance was calculated by unpaired t-test; ***, P< 0.0001; **, P < 0.001.

Consistent with these data, we found that when we quantified vaginal epithelial cells with a Clue Cell-like appearance (i.e., covered with bound bacteria)^15^ in digests of the epithelium in Vagina Chip containing the dysbiotic BVC1 consortium, we observed a decrease in the number of these cells in the presence of cervical mucus (**Fig. 2C,D**). Not surprisingly, this reduction in bacterial cell number induced by the presence of cervical mucus was also accompanied by a concomitant increase in vaginal epithelial cell viability (retained cell number) (**Fig. 2E**), as well as significant downregulation of the proinflammatory cytokines, IL-8, IL-10, Rantes (CCL5), TNF-α, MIP-1β and IL-1α after 72 hours of co-culture (**Fig. 3**). These results demonstrate that cervix chip mucus can directly influence the epithelium to dampen production of inflammatory cytokines and this correlates with protection of the epithelium against injury.

**Fig. 3.**
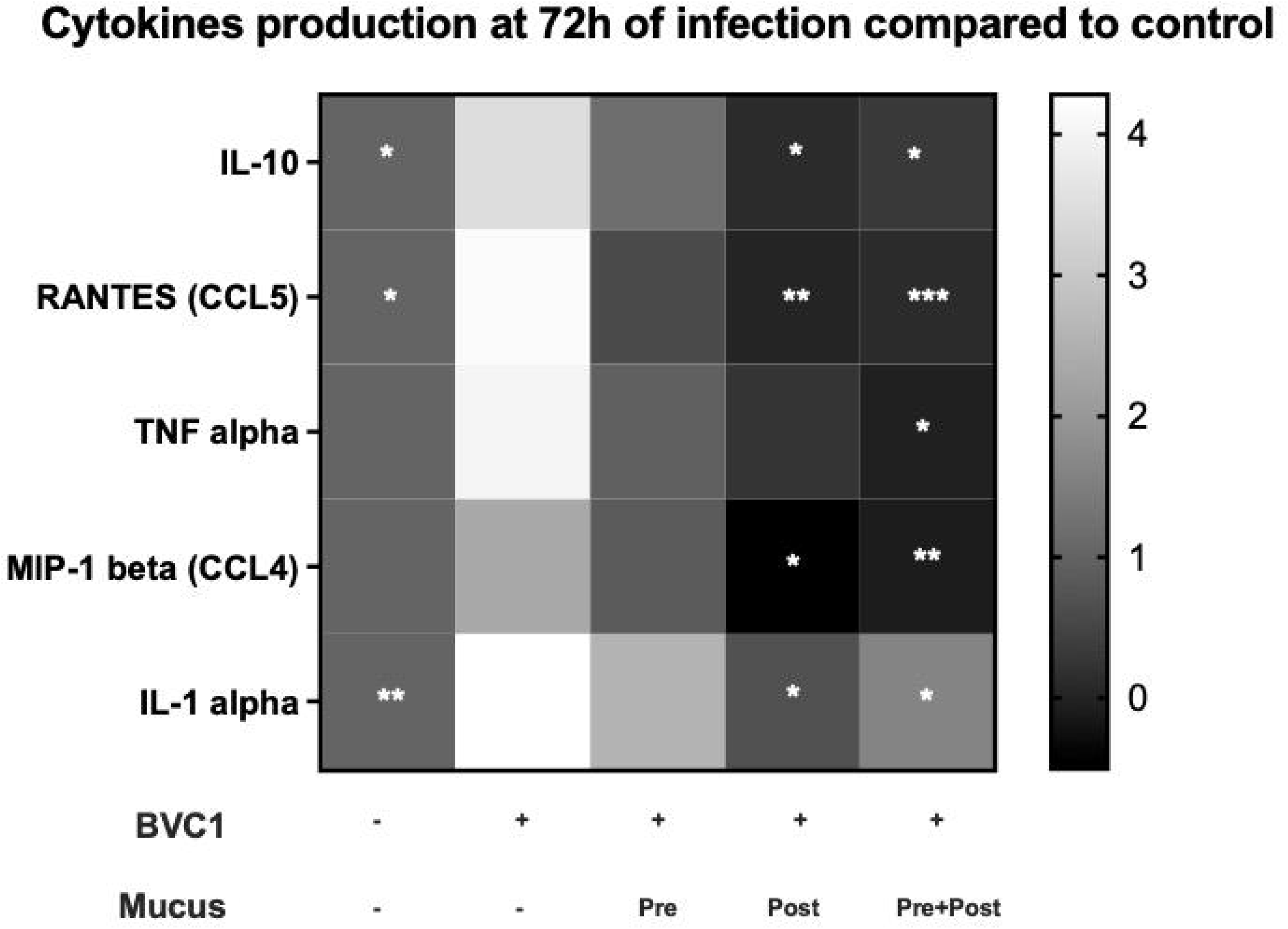
Cytokine Production by the Vagina Chip. Heat map showing the innate immune response of vaginal epithelium cultured on-chip for 72 hours in the absence (Control) or presence of BVC1 consortium perfused either without or with mucus-containing effluents from Cervix Chips that were added 1 day prior to addition of bacteria (Pre), 1 day after BVC 1 addition (Post), or continuously for the entire 3 day culture starting 1 day prior to addition of bacteria (Pre+Post). IL-10, RANTES(CCL5), TNF-α, MIP-1β, IL-1α protein levels in the epithelial channel effluents were normalized for cell number. The gray scale represents fold change in cytokine levels relative to levels in the control chip. n=4-10 individual chips for each group from 4 independent experiments; significance was calculated by unpaired t-test; *P<0.05, ***P < 0.001, ****P < 0.0001 compered to BVC1.

### Suppression of growth of G. vaginalis in Vagina Chip Effluents

To explore whether the cervical mucus acts directly to suppress bacterial cell growth or indirectly by altering vaginal cell physiology, we next compared the growth of *G. vaginalis* in mucus-containing effluent samples collected from the epithelial channel of control Cervix Chips (perfused with HBSS) versus effluents from Vagina Chips that were perfused with Cervix Chip derived mucus-containing effluent for 1 day, with the bacteria cultured directly in the Hank’s balanced salt solution (HBBS) that is used to perfuse the apical channels of our chips as a control. Our results demonstrate that *G. vaginalis* bacteria grew well in the Cervix Chip mucus in 2D culture, but growth was suppressed when they were cultured in effluents from the Vagina Chip perfused with similar Cervix Chip derived mucus-containing effluent or in HBSS that lacks critical (**Fig. 4A,B**). Importantly, when similar studies were carried out after addition of 50% bacterial broth to provide optimal nutrient conditions, bacterial growth was restored in the control HBSS sample, but not in the sample from the Vagina Chip exposed to Cervix Chip-derived mucus effluent (**Fig. 4C,D**). These findings suggest that mucus components produced by the Cervix Chip induce the cells lining the Vagina Chip to elaborate factors that suppress *G. vaginalis* growth.

**Fig. 4.**
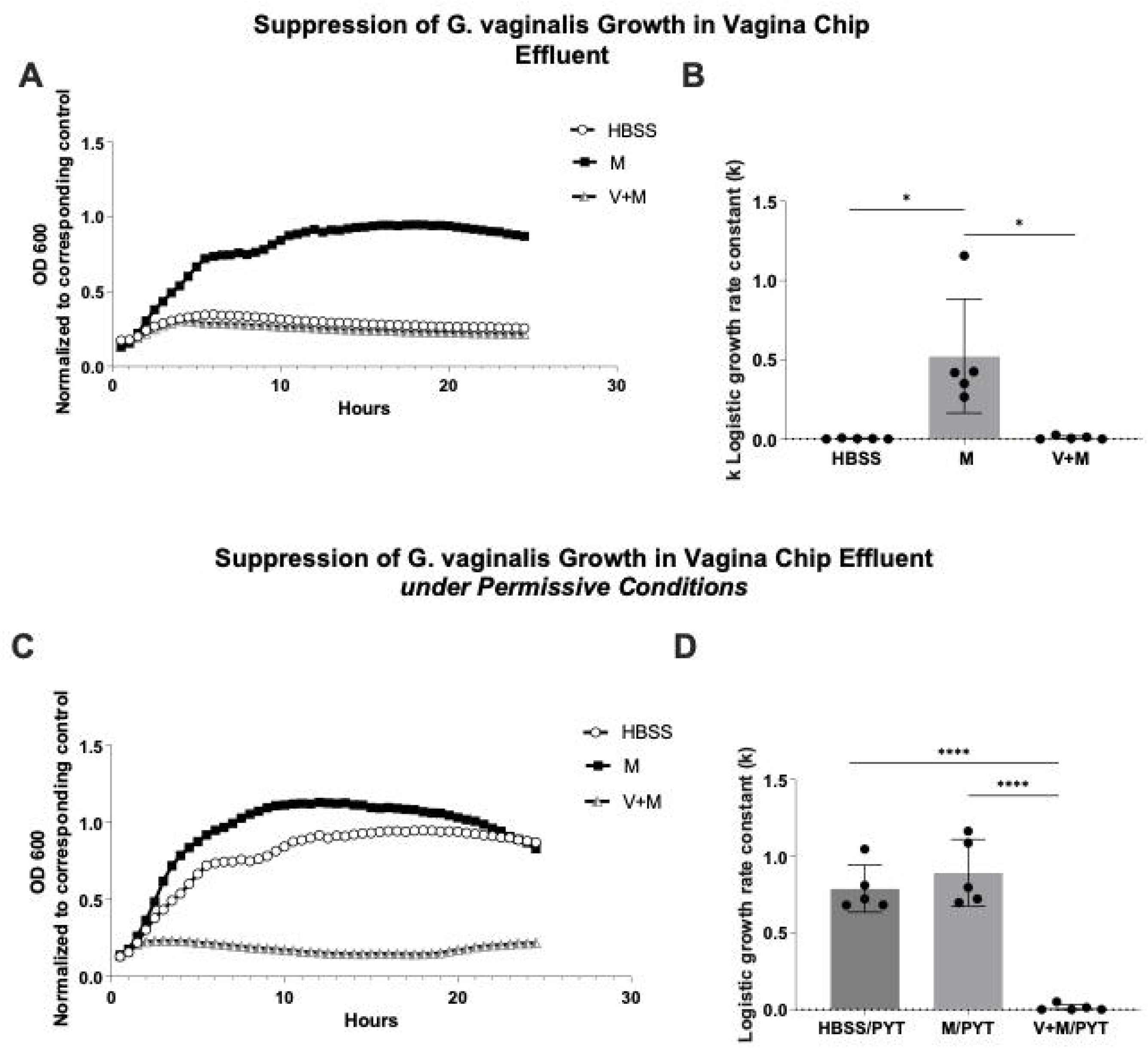
*Suppression of G. vaginalis growth in Vagina Chip effluents* in 2D culture. **A)** *G. vaginalis* growth in conventional culture wells measured by recording optical density (OD) measurements taken every 30 minutes for 24 hours when the bacteria were placed within mucus-containing Cervix Chip effluent (M), similar effluent perfused through a Vagina Chip for 1 day (V+M), or an HBSS control. **B)** Logistic growth rate constant (k) for bacterial growth in panel A. **C)** Bacterial growth in permissive conditions in which the effluents shown in A were supplemented with 50% bacterial broth (PYT). Note that while permissive conditions allowed for growth in the HBSS control group, *G. vaginalis* growth was still suppressed in the V+M group. **D)** Logistic growth rate constant (k) for bacterial growth in panel C n=5; *P<0.05, ***P < 0.001, ****P < 0.0001.

### Cervix Chip mucus alters the vaginal secretome

To further explore the effects of the cervical mucus on the vaginal epithelium, we conducted mass spectrometry analysis to compare the proteome composition of the Cervix Chip effluent before or after exposure to the Vagina Chip versus the untreated Vagina Chip effluent. Out of the 1752 proteins identified (**Supplementary Table 1**), 103 were found to be differentially abundant as determined by fold change (|log2 fc| >= 1), p_adj_ ≤ 0.05) in Cervix Chip effluents that had passed through the Vagina Chip versus either the Cervix Chip efffluent or Vagina Chip effluent alone. Significant changes in expression of multiple proteins were observed, with 64 proteins showing increased expression (**Fig. 5A**) and 39 proteins decreasing (**Fig. 5B**), with the most noteworthy alterations in protein expression highlighted in a volcano plot (**Fig. 5C**). PCA analysis of the proteomics data also revealed distinct segregation among these sample groups, indicating notable variances in protein expression in effluents from Vagina Chips exposed to Cervix Chip mucus compared to those from untreated Cervix or Vagina Chips alone (**Fig. 5D**).

**Fig. 5.**
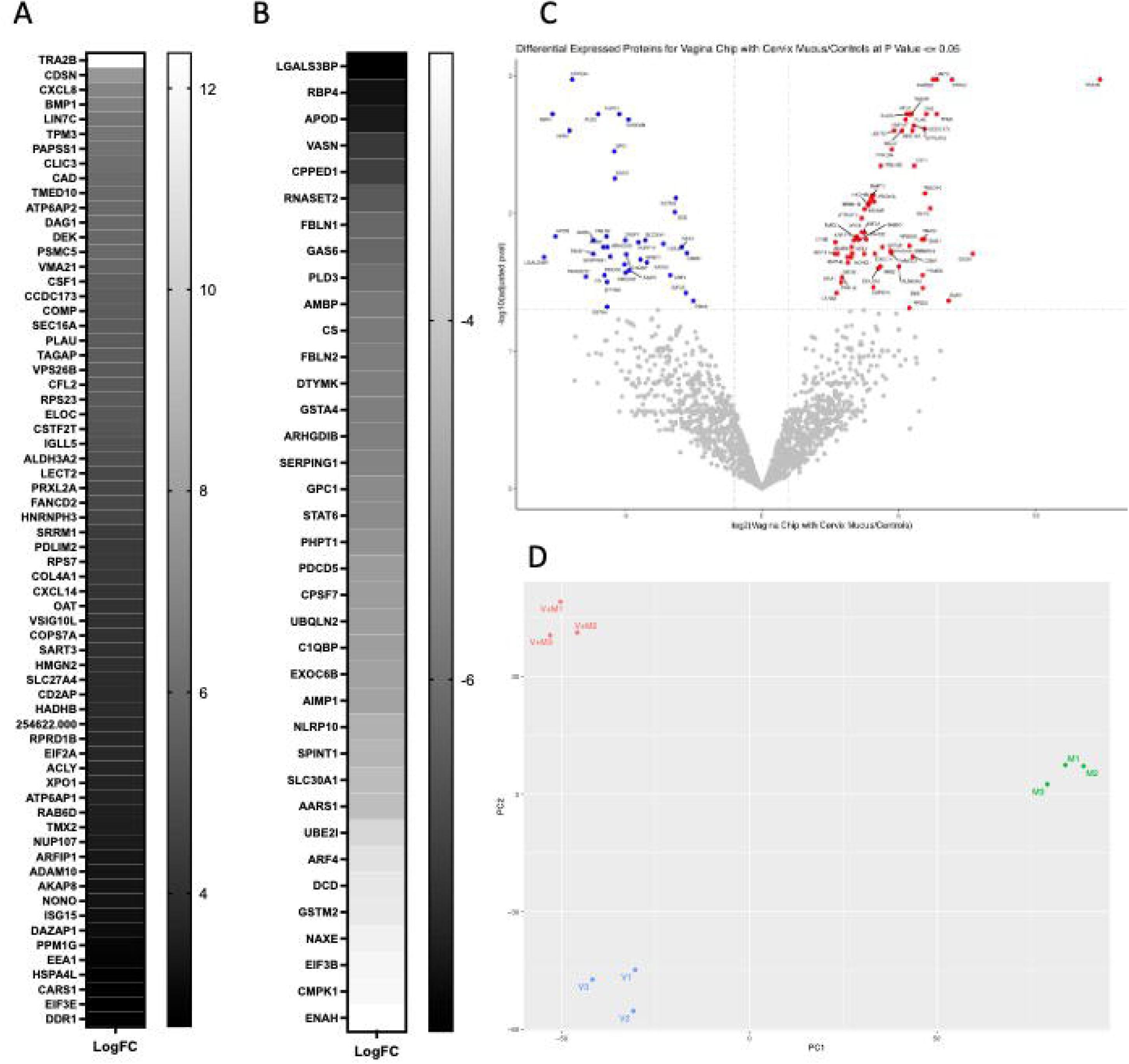
Cervical mucus alters the vaginal secretome. Mass spectrometry analysis of the Vagina Chip samples effluents pre-exposed to Cervixc Chip mucus-containing effluent for 1 day (V+M) compared to those found within the Vagina Chip effluent (V) and Cervical Chip mucus-containing effluent (M) alone. **A)** Up regulated and **B)** Downregulated proteins in V+M compared V and M, as determined by fold change (|log2 fc| >= 1, p_adj_ ≤ 0.05) **C**) Volcano plot showing differentially expressed proteins in V+M comperd to V and M. The plot was constructed using the normalized protein expression data with the negative logarithm of the adjusted p-value represented on the y-axis and the log2 fold change represented on the x-axis. Each dot on the plot corresponds to a protein, with color-coding used to indicate the statistical significance of differential expression. Proteins with a *p* value < 0.05 and fold change > 2 were colored red, while proteins with a *p* value < 0.05 and fold change < -2 were colored blue (Gray, proteins with *p* value > 0.05).

Interestingly, using STRING analysis, which incorporates both physical protein-protein interactions and functional associations from various sources (e.g., automated text mining, computational interaction predictions from co-expression, conserved genomic context, databases of interaction experiments, and curated sources of known complexes/pathways),^16^ we found that 3 of the 37 down-regulated proteins exhibit calcium channel inhibitor activity (PHPT1,AMBP,SLC30A1). Previous research has shown that G*. vaginalis* strongly induces epithelial calcium influx and contraction.^17^ In addition, 6 of the down-regulated proteins are ECM molecules (LGALS3BP, GPC1, AMBP, SERPING1, VASN, FBLN1, FBLN2), which may play a role in *G. vaginalis* adhesion and biofilm formation.^18^ Finally, 3 down-regulated proteins were members of the Lipocalin family (AMBP, APOD, RBP4), which is known for its role in regulating inflammation and antioxidant responses.^19^

The STRING analysis additionally showed that 17 of the up-regulated proteins are RNA binding proteins (RBPs). Previous studies have highlighted the vital role of RBPs bacterial replication by binding to and regulating their RNAs.^20^ These proteins also play a crucial role in the immune system’s response to viral infections by regulating viral RNA stability and translation.^21^ Considering that BV increases susceptibility to sexually transmitted infections, including viral infections, these findings further support the potential involvement of RBPs in the immune response within the reproductive tract. Twenty five proteins related to the male reproductive system also were found to be upregulated. This finding is significant because past studies have established a notable link between BV and infertility.^6^ For instance, one of the proteins identified, CSTF2T, has the potential to contribute to sperm adhesion to the zona pellucida^22^ while the TMED10 protein may be involved in in sperm capacitation and the acrosome reaction.^23^

Importantly, exposure of the Vagina Chip also resulted in enhanced production of potential antimicrobial proteins PLAU and WASF2. PLAU is a serine protease with immunomodulatory functions^24^ while WASF2 is a member of the Wiskott-Aldrich syndrome protein family that regulates autophagy and inflammasome activity.^25^ One of the prominent down-regulated proteins, GNS, is an N-acetylglucosamine-6-sulfatase. This is interesting because BV is often associated with the breakdown of mucins, which is necessary for these dysbiotic bacteria to colonize the vagina.^26^ Thus, downregulation of GNS could contribute to the inhibition of growth of the dysbiotic bacteria we observed by increasing glycoprotein sulfation and thereby, preventing mucin degradation.

### Potential role of exosomes as mediators of the effects of cervical mucus on the Vagina Chip

Exosomes, which are small extracellular vesicles containing nucleic acids, lipids, and proteins, play a significant role in intercellular communication in the female reproductive tract by modulating the immune system and promoting tissue repair.^27^ This is accomplished by presenting antigenic peptides, regulating gene expression through exosomal miRNA, and inducing different signaling through exosomal surface ligands. Importantly, when we carried out STRING analysis of the proteins differentially express in Vagina Chip effluents exposed to Cervix Chip mucus, we found that a significant proportion of the differentially expressed proteins were associated with exosomes. Specifically, 23 out of 37 down-regulated proteins and 17 out of 64 upregulated proteins were found to be linked to extracellular exosomes (**Supplementary Table 2**), Notable among the upregulated proteins were DDR1^28^ and COMP,^29^ which regulate cellular adhesion to the ECM and its remodeling, which influence bacterial adhesion.^30^ Conversely, among the down-regulated proteins, five ECM proteins (AMBP, FBLN1, GPC1, LGALS3BP, and SERPING1) were identified, which may also influence bacterial adhesion. Interestingly, SERPING1 functions as a regulator of the complement system^31^ and three of the down-regulated proteins (AMBP, SERPING1, and SPINT1) belong to the Kunitz family of serine protease inhibitors that are involved in coordinating inflammation.^32^

### Cervicovaginal antimicrobial peptides

Additionally, we identified 12 antimicrobial peptides that were present in the Cervix and Vagina Chip effluents (**Table 1**). Of these, 6 (Dermcidin, Ubiquicidin, Chemerin, Acipensin 6, hSAA1, and Psoriasin) were present in both Vagina and Cervix Chip effluents, 1 was solely produced by the Vagina Chip (KAMP-19), and 5 were exclusively produced by the Cervix Chip. No antimicrobial peptides were specifically induced in Vagina Chips exposed to Cervix Chip effluents. The Cervix Chip-derived antimicrobial peptides include Histone H4, Histone H3, CXCL1, BHP and Chromacin. Histones and their fragments have a variety of antimicrobial actions and functions, including bacterial cell membrane permeabilization, penetration into the membrane followed by binding to bacterial DNA and/or RNA, binding to bacterial lipopolysaccharide (LPS) and neutralizing its toxicity, and entrapping pathogens as a component of neutrophil extracellular traps.^33, 34^ It is noteworthy that *P. bivia*, which is included in our bacterial consortium, has been shown to produce high concentrations of LPS.^35^ CXCL1 also inhibits growth of *E. coli* and *S. aureus* in vitro^36^ and BHP impedes the growth of *M. luteus, S. epidermidis*, and several fungi (e.g.,*C. albicans, S. cerevisiae, and A. nidulans)*^37^, while Chromacin suppresses the growth of *Bacillus megaterium* and *Micrococcus luteus*.^38^

**Table 1.**
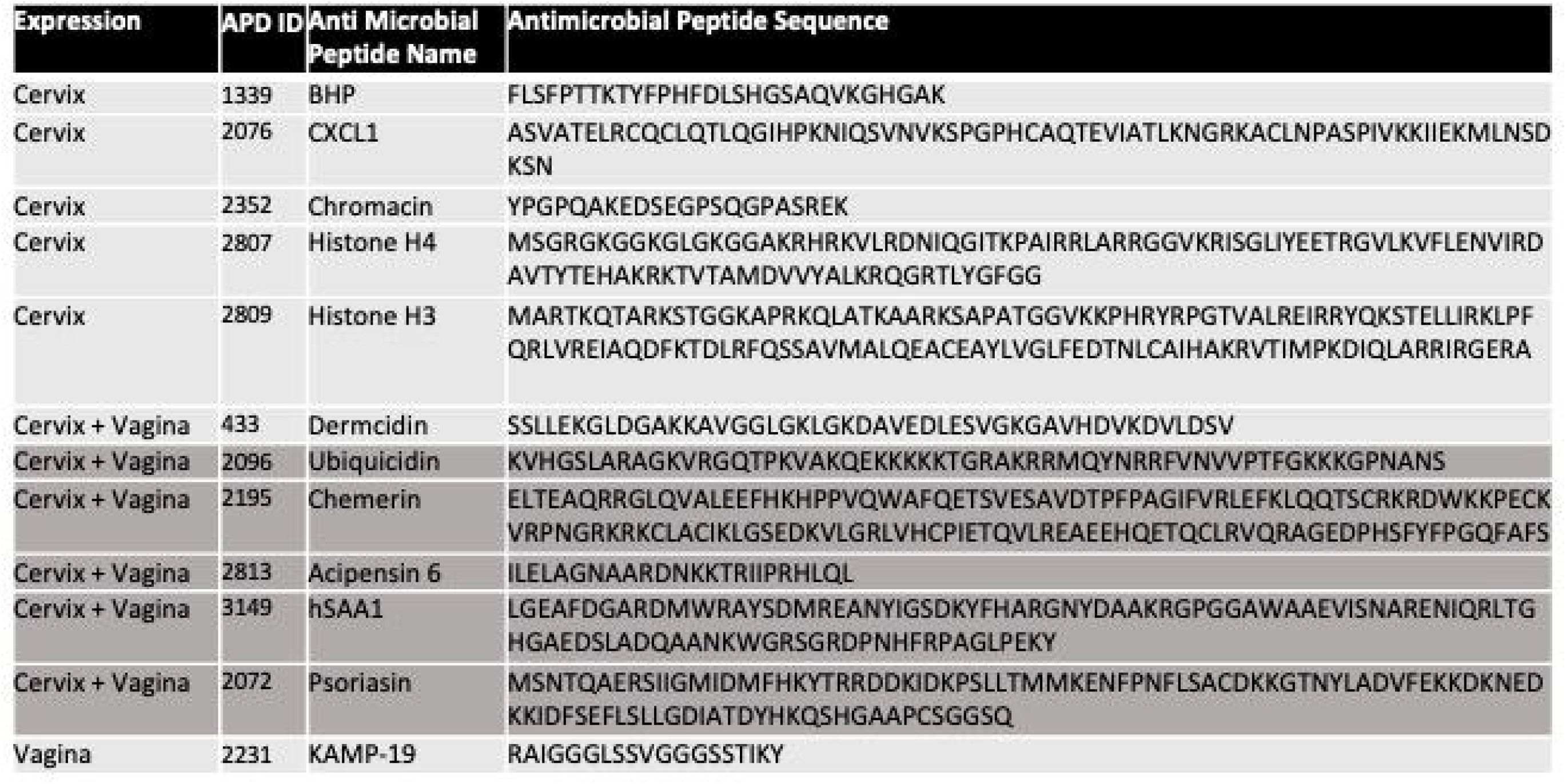

### Comment

#### Principal Findings

These data show that mucus containing effluents from human Cervix Chips suppress growth of dysbiotic microbiota, associated inflammation and epithelial cell injury in human Vagina Chips. By analyzing the differentially abundant proteins in the secretome of Vagina Chips following treatment with the Cervix chip effluents, we identified multiple proteins that may contribute to this protective response and that potentially could be used as clinical biomarkers for monitoring health of the female reproductive tract in the future.

#### Results in the Context of What is Known

Maintaining homeostasis is crucial for the health of epithelial barriers, which can be disrupted during infections or injury. Inflammation plays a vital role in supporting the body’s defense against pathogens, promoting tissue healing, and restoring homeostasis.^39^ However, chronically high levels of proinflammatory cytokines that undermine normal protective immune signals have been linked to an imbalanced microbiome and compromised epithelial cell stability.^40^ This study presents evidence that communication between cervical and vaginal tissues in the lower reproductive tract via transfer of cervical mucus-containing secretions helps to suppress vaginal inflammation in the presence of a dysbiotic microbiome. There is a growing body of clinical evidence suggesting that cervical procedures may disrupt this crucial communication between cervical and vaginal epithelium, and lead to changes in the composition of vaginal microbiome.^41, 42^ Our results support this observation and suggest that it is a direct effect of reducing cervical mucus transfer to the vagina, which could only be studied directly using this type of engineered in vitro model.

The recurrence of abnormal vaginal flora after the treatment of BV (e.g., with metronidazole) is commonly detected in most women,^8^ however, the underlying factors contributing to these recurrences remain elusive. Our findings suggest that alterations in cervical mucus levels may influence the susceptibility of the vaginal epithelium to BV infection. Therefore, an imbalance in the cervicovaginal mucus may be a possible contributing factor to the high rate of BV recurrence. In this context, it is important to note that we identified 5 cervical antimicrobial peptides that appear to play a role in the antimicrobial effects we observed on-chip. These findings suggest that interactions between antimicrobial peptides and the host vaginal epithelium can enhance innate immune protection against dysbiotic flora.

Immune effectors and specialized stromal cells at epithelial surfaces produce cytokines and antimicrobial defenses to orchestrate tissue repair and to minimize opportunistic infections. Exosomes can act as mediators for this form of inter-tissue communication. We identified 40 exosomal proteins produced by vaginal epithelium that were modulated by exposure to cervical mucus produced in the human Cervix Chip. Human cervicovaginal exosomes have been previously shown to be part of the female innate defense system and to protect against HIV-1 infection^43^ as well as bacterial toxins.^44^ Exosomes are also being actively explored as potential therapeutic agents and drug delivery vehicles. Thus, the ability to study the role of exosomes in host-microbiome interactions in the female reproductive tract in vitro using the human Organ Chip models described here may facilitate development of novel treatments for vaginal dysbiosis as well as other diseases of the female reproductive tract.

#### Clinical Implications

Our study has important clinical implications as it has the potential to identify new targets for diagnosis and treatment of vaginal diseases. Identifying patients with a high likelihood of recurrent vaginal dysbiosis can help to customize their treatment plan and prevent complications. In this study, we identified multiple proteins and antimicrobial peptides that may contribute to the protective response against dysbiotic microbiota and associated inflammation and injury to the vaginal epithelium. These proteins and peptides could potentially be used as clinical biomarkers for monitoring the health of the female reproductive tract in the future. Several proteins we identified (e.g., TPM3, PLAU, ALDH3A2, GAS6, DTYMK, SERPING, STAT6, CMPK1) are known to be targeted by existing approved drugs (Progesterone, Urokinase, Disulfiram, Warfarin, Zidovudine, Rhucin, Indomethacin, and Gemcitabine, respectively). Thus, if these molecule actively contribute to the BV disease phenotype, one or more these therapeutics could be added to current clinical regimens.

#### Research Implications

Our results show the value of human Organ Chip technology for studying vaginal health and diseases of the female reproductive tract. However, further research is needed to evaluate the effects of these compounds as well as modulators of the other putative targets we identified on maintaining vaginal homeostasis and a healthy microbiome.

#### Strengths and Limitations

While the human Vagina and Cervix Chips used in this study replicate many physiological and patholphysiological features of the female reproductive tract, we did not incorporate immune cells in models. As these cells play a crucial role in mounting antibacterial immune responses, the model would be strengthened by incorporating them in the future. Additionally, it should be noted that the Organ Chips we used were created with epithelial cells from a single human donor and thus, these studies should be extended to include chips lined by cells from multiple donors from different ethnic groups in the future.

#### Conclusions

This study highlights the crucial role that cervical mucus plays in maintaining vaginal health and preventing dysbiosis-related complications. Our results directly demonstrate that cervical mucus-containing secretions can suppress the growth of dysbiotic microbiota as well as associated inflammation and epithelial cell injury in the human Vagina Chip. We also identified several proteins and antimicrobial peptides that could serve as clinical biomarkers for monitoring the health of the female reproductive tract and potentially be targeted for treatment of vaginal dysbiosis. The study also sheds light on the potential role of exosomes in inter-tissue communication and immune protection against dysbiotic flora in the female reproductive tract. In addition, these findings could have important clinical implications, particularly for identifying patients with a high risk of recurrent dysbiosis and customizing their treatment plan. Further research is needed to evaluate the effects of modulating the potential molecular mediators we identified on maintaining healthy vaginal microbial homeostasis. However, these findings provide further evidence showing that human Organ Chip models can provide a valuable tool for studying host-microbiome interactions in the female reproductive tract as well as for identifying potential clinical biomarkers and therapeutic targets for patients with vaginal dysbiosis and other related diseases.

## ACKNOWLEDGMENTS

This research was sponsored by the funding from the Bill and Melinda Gates Foundation (OPP1173198 & INV-035977 to D.E.I., OPP1189217 to J.R. and INV-031642 to S.R-N.) and the Wyss Institute for Biologically Inspired Engineering (D.E.I.).

## AUTHORS CONTRIBUTIONS

O.G.: conceptualization, data curation, formal analysis, investigation, writing - original draft. A.G.: investigation, writing - review & editing. Z.I.: conceptualization, writing - review & editing. A.S.: conceptualization. H.R.: investigation.J.C.: investigation. B.B.: investigation, proteomics methodology. S.S.: software. G.G.: supervision, writing - review & editing. A.J.: supervision, writing - review & editing. D.E.I.: conceptualization, supervision, funding acquisition, writing - review & editing.

## Condensation page

### Tweetable statement

Human Organ Chips reveal that cervical mucus plays a critical role in preventing complications from vaginal dysbiosis. The work led to identification of potential diagnostic biomarkers and therapeutic targets for managing bacterial vaginosis.

## AJOG at a Glance

A. Why was this study conducted?
  - To assess the role of cervical mucus in preventing vaginal dysbiosis-related complications
B. What are the key findings?
  - Cervical mucus protects the vaginal epithelium from inflammation and epithelial cell injury caused by a dysbiotic microbiome
  - Proteomic analysis of proteins produced by the Vagina Chip following treatment with Cervix Chip mucus revealed potential diagnostic biomarkers and therapeutic targets for managing bacterial vaginosis.
C. What does this study add to what is already known?
  - The study highlights the potential significance of cervical mucus in human vaginal physiology and pathophysiology
  - The study also demonstrates the potential value of Organ Chip technology for studies focused on health and disease of the female reproductive tract

## REFERENCES

1. Lacroix G, Gouyer V, Gottrand F, Desseyn JL. The Cervicovaginal Mucus Barrier. Int J Mol Sci 2020;21.

2. Vagios S, Mitchell CM. Mutual Preservation: A Review of Interactions Between Cervicovaginal Mucus and Microbiota. Front Cell Infect Microbiol 2021;11:676114.

3. McLoughlin K, Schluter J, Rakoff-Nahoum S, Smith AL, Foster KR. Host Selection of Microbiota via Differential Adhesion. Cell Host Microbe 2016;19:550–9.

4. Juliana NCA, Suiters MJM, Al-Nasiry S, Morre SA, Peters RPH, Ambrosino E. The Association Between Vaginal Microbiota Dysbiosis, Bacterial Vaginosis, and Aerobic Vaginitis, and Adverse Pregnancy Outcomes of Women Living in Sub-Saharan Africa: A Systematic Review. Front Public Health 2020;8:567885.

5. Janulaitiene M, Paliulyte V, Grinceviciene S, et al. Prevalence and distribution of Gardnerella vaginalis subgroups in women with and without bacterial vaginosis. BMC Infect Dis 2017;17:394.

6. Ravel J, Moreno I, Simon C. Bacterial vaginosis and its association with infertility, endometritis, and pelvic inflammatory disease. Am J Obstet Gynecol 2021;224:251–57.

7. Van De Wijgert J, Jespers V. The global health impact of vaginal dysbiosis. Res Microbiol 2017;168:859–64.

8. Neal CM, Kus LH, Eckert LO, Peipert JF. Noncandidal vaginitis: a comprehensive approach to diagnosis and management. Am J Obstet Gynecol 2020;222:114–22.

9. Mahajan G, Doherty E, TO T, et al. Vaginal microbiome-host interactions modeled in a human vagina-on-a-chip. Microbiome 2022;10:201.

10. Izadifar Z, Cotton J, Chen C, et al. Mucus production, host-microbiome interactions, hormone sensitivity, and innate immune responses modeled in human endo- and ecto-cervix chips. bioRxiv 2023:2023.02.22.529436.

11. Ravel J, Gajer P, Abdo Z, et al. Vaginal microbiome of reproductive-age women. Proc Natl Acad Sci U S A 2011;108 Suppl 1:4680–7.

12. Ravel J, Brotman RM, Gajer P, et al. Daily temporal dynamics of vaginal microbiota before, during and after episodes of bacterial vaginosis. Microbiome 2013;1:29.

13. Bloom SM, Mafunda NA, Woolston BM, et al. Cysteine dependence of Lactobacillus iners is a potential therapeutic target for vaginal microbiota modulation. Nat Microbiol 2022;7:434–50.

14. Kall L, Storey JD, Noble WS. Non-parametric estimation of posterior error probabilities associated with peptides identified by tandem mass spectrometry. Bioinformatics 2008;24:i42–8.

15. Gardner HL, Dukes CD. Haemophilus vaginalis vaginitis: a newly defined specific infection previously classified non-specific vaginitis. Am J Obstet Gynecol 1955;69:962–76.

16. Szklarczyk D, Kirsch R, Koutrouli M, et al. The STRING database in 2023: protein-protein association networks and functional enrichment analyses for any sequenced genome of interest. Nucleic Acids Res 2023;51:D638–D46.

17. Abbasian B, Shair A, O’Gorman DB, et al. Potential Role of Extracellular ATP Released by Bacteria in Bladder Infection and Contractility. mSphere 2019;4.

18. Hardy L, Cerca N, Jespers V, Vaneechoutte M, Crucitti T. Bacterial biofilms in the vagina. Res Microbiol 2017;168:865–74.

19. Rassart E, Desmarais F, Najyb O, Bergeron KF, Mounier C. Apolipoprotein D. Gene 2020;756:144874.

20. Van Assche E, Van Puyvelde S, Vanderleyden J, Steenackers HP. RNA-binding proteins involved in post-transcriptional regulation in bacteria. Front Microbiol 2015;6:141.

21. Gao Q, Jiang M, Zhao Y, et al. eIF4A3 Promotes RNA Viruses’ Replication by Inhibiting Innate Immune Responses. J Virol 2022;96:e0151322.

22. Tardif S, Akrofi AS, Dass B, Hardy DM, Macdonald CC. Infertility with impaired zona pellucida adhesion of spermatozoa from mice lacking TauCstF-64. Biol Reprod 2010;83:464–72.

23. Castillo J, Bogle OA, Jodar M, et al. Proteomic Changes in Human Sperm During Sequential in vitro Capacitation and Acrosome Reaction. Front Cell Dev Biol 2019;7:295.

24. Jin T, Bokarewa M, Tarkowski A. Urokinase-type plasminogen activator, an endogenous antibiotic. J Infect Dis 2005;192:429–37.

25. Lee PP, Lobato-MARQUEZ D, Pramanik N, et al. Wiskott-Aldrich syndrome protein regulates autophagy and inflammasome activity in innate immune cells. Nat Commun 2017;8:1576.

26. Roberton AM, Wiggins R, Horner PJ, et al. A novel bacterial mucinase, glycosulfatase, is associated with bacterial vaginosis. J Clin Microbiol 2005;43:5504–8.

27. Kalluri R, Lebleu VS. The biology, function, and biomedical applications of exosomes. Science 2020;367.

28. Duan X, Xu X, Zhang Y, Gao Y, Zhou J, Li J. DDR1 functions as an immune negative factor in colorectal cancer by regulating tumor-infiltrating T cells through IL-18. Cancer Sci 2022;113:3672–85.

29. Posey KL, Coustry F, Hecht JT. Cartilage oligomeric matrix protein: COMPopathies and beyond. Matrix Biol 2018;71–72:161-73.

30. Marrs CN, Knobel SM, Zhu WQ, Sweet SD, Chaudhry AR, Alcendor DJ. Evidence for Gardnerella vaginalis uptake and internalization by squamous vaginal epithelial cells: implications for the pathogenesis of bacterial vaginosis. Microbes Infect 2012;14:500–8.

31. Zeerleder S. C1-inhibitor: more than a serine protease inhibitor. Semin Thromb Hemost 2011;37:362–74.

32. Shigetomi H, Onogi A, Kajiwara H, et al. Anti-inflammatory actions of serine protease inhibitors containing the Kunitz domain. Inflamm Res 2010;59:679–87.

33. Wira CR, Patel MV, Ghosh M, Mukura L, Fahey JV. Innate immunity in the human female reproductive tract: endocrine regulation of endogenous antimicrobial protection against HIV and other sexually transmitted infections. Am J Reprod Immunol 2011;65:196–211.

34. Drab T, Kracmerova J, Hanzlikova E, et al. The antimicrobial action of histones in the reproductive tract of cow. Biochem Biophys Res Commun 2014;443:987–90.

35. Aroutcheva A, Ling Z, Faro S. Prevotella bivia as a source of lipopolysaccharide in the vagina. Anaerobe 2008;14:256–60.

36. Yang D, Chen Q, Hoover DM, et al. Many chemokines including CCL20/MIP-3alpha display antimicrobial activity. J Leukoc Biol 2003;74:448–55.

37. Conlon JM, Kolodziejek J, Nowotny N. Antimicrobial peptides from the skins of North American frogs. Biochim Biophys Acta 2009;1788:1556–63.

38. Strub JM, Goumon Y, Lugardon K, et al. Antibacterial activity of glycosylated and phosphorylated chromogranin A-derived peptide 173-194 from bovine adrenal medullary chromaffin granules. J Biol Chem 1996;271:28533–40.

39. Rathinam VAK, Chan FK. Inflammasome, Inflammation, and Tissue Homeostasis. Trends Mol Med 2018;24:304–18.

40. Clavel T, Haller D. Bacteria- and host-derived mechanisms to control intestinal epithelial cell homeostasis: implications for chronic inflammation. Inflamm Bowel Dis 2007;13:1153–64.

41. Kawahara R, Fujii T, Kukimoto I, et al. Changes to the cervicovaginal microbiota and cervical cytokine profile following surgery for cervical intraepithelial neoplasia. Sci Rep 2021;11:2156.

42. Wiik J, Sengpiel V, Kyrgiou M, et al. Cervical microbiota in women with cervical intra-epithelial neoplasia, prior to and after local excisional treatment, a Norwegian cohort study. BMC Womens Health 2019;19:30.

43. Smith JA, Daniel R. Human vaginal fluid contains exosomes that have an inhibitory effect on an early step of the HIV-1 life cycle. AIDS 2016;30:2611–16.

44. Keller MD, Ching KL, Liang FX, et al. Decoy exosomes provide protection against bacterial toxins. Nature 2020;579:260–64.

